# Antimicrobial resistance and virulence characteristics of *Klebsiella pneumoniae* isolates in Kenya

**DOI:** 10.1101/2022.02.01.478614

**Authors:** Angela W. Muraya, Cecilia Kyany’a, Shahiid Kiyaga, Hunter J. Smith, Caleb Kibet, Melissa J. Martin, Josephine Kimani, Lillian Musila

## Abstract

*Klebsiella pneumoniae* is a globally significant opportunistic pathogen causing healthcare-associated and community-acquired infections. This study examined the epidemiology and the distribution of resistance and virulence genes in clinical *K. pneumoniae* strains in Kenya. Eighty-nine *K. pneumoniae* isolates were collected over six years from five counties in Kenya and were analyzed using whole genome sequencing and bioinformatics. These isolates were obtained from community-acquired (62/89) and healthcare-associated infections (21/89), and the hospital environment (6/89). Genetic analysis revealed the presence of *bla_NDM-1_* and *bla_OXA-181_* carbapenemase genes and the *armA* and *rmtF* genes known to confer pan-aminoglycoside resistance. The most abundant extended-spectrum beta-lactamase genes identified were *bla_CTX-M-15_* (36/89), *bla_TEM_* (35/89), and *bla_OXA_* (18/89). In addition, one isolate had a mobile colistin resistance gene (*mcr-8*). Fluoroquinolone resistance-conferring mutations in *gyrA* and *parC* genes were also observed. The most notable virulence factors were those associated with hyper-virulence (*rmpA/A2* and *magA*), yersiniabactin (*ybt*), salmochelin (*iro*), and aerobactin (*iuc* and *iutA*). Thirty-eight distinct sequence types were identified, including known global lineages ST14, ST15, ST147, and ST307, and a regional clone ST17 implicated in regional outbreaks. In addition, this study genetically characterized two potential hypervirulent isolates and two community-acquired ST147 high-risk clones that contained carbapenemase genes, yersiniabactin, and other multidrug resistance genes. These results demonstrate that the resistome and virulome of Kenyan clinical and hospital environmental *K. pneumoniae* isolates are diverse. The reservoir of high risk-clones capable of spreading resistance and virulence factors have the potential to cause unmanageable infection outbreaks with high morbidity and mortality.

**AUTHOR SUMMARY:** *Klebsiella pneumoniae* is one of the human-disease-causing bacteria that easily acquires and spreads antibiotic resistance genes and is thus a serious threat to human health. We studied both the antibiotic resistance genes and the genes it uses to cause disease (virulence). Forty-two percent of our isolates were multidrug resistant (MDR). They carried several resistance and virulence genes bound in mobile circular DNA molecules called plasmids which easily migrate and spread the genes between bacteria. We identified 38 distinct *K. pneumoniae* strains (STs) distributed within the study sites. Fifteen isolates were classified under the groups of *K. pneumoniae* strains known to cause global infection outbreaks such as ST14, ST15 and ST147 collected from Nairobi and Kisumu, hotspot areas for spread of resistance. In particular, two ST147 isolates were resistant to carbapenems and one isolate to colistin, which are last line antibiotics. We also identified two isolates with the potential to cause high levels of disease. We concluded that the presence of highly resistant and virulent strains in the hospital and community demonstrates a need for the continuous monitoring and management of MDR *K. pneumoniae* infections to prevent disease outbreaks that are difficult to control and that lead to high death rate.

## INTRODUCTION

*Klebsiella pneumoniae* (KP) is a gram-negative, rod-shaped ubiquitous bacterium that inhabits soil, water, and sewage ecosystems. It is also found on various human body sites and organ systems, including skin, nose, throat, and intestinal tract, as part of the natural microflora (1). *K. pneumoniae* is a prominent member of the *Klebsiella pneumoniae* species complex (KpSC), consisting of four species commonly associated with human infections, including *Klebsiella pneumoniae subsp. pneumoniae, Klebsiella quasipneumoniae subsp. quasipneumoniae, Klebsiella quasipneumoniae subsp. similipneumoniae* and *Klebsiella variicola subsp. variicola* (2). These species cause infections such as pneumonia, urinary tract infections, soft tissue and wound infections, septicemia, and pyogenic liver abscesses (3).

The World Health Organization (WHO) declared antimicrobial resistance (AMR) as one of the top 10 most serious global public health threats facing humanity (4). The WHO lists *K. pneumoniae* as one of the AMR bacteria of concern due to its demonstrated proclivity for developing antimicrobial resistance to many classes of antibiotics such as penicillins, cephalosporins, and quinolones (5–7) which are typically used to treat KP infections. This resistance is due to both chromosomal-encoded and plasmid-encoded genes. The abundance of AMR genes carried on plasmids and mobile genetic elements have earned KP its reputation as a “key trafficker” of AMR genes between Klebsiella species and other Enterobacterales, illustrating its importance to AMR spread and development (8). As carbapenems are considered one of the last resort treatments for multidrug-resistant (MDR) KP, i.e., isolates resistant to 3 or more drug classes (9), the global increase in carbapenem resistance (10) presents a threat to public health. The situation is aggravated by the isolation of colistin-resistant KP (11) since colistin is used as a last-line antibiotic for treating carbapenem-resistant KP. Global MDR high-risk lineages such as ST14, ST15, ST147, ST307, and ST607 have been identified and have spread rapidly across the globe increasing the need for global AMR surveillance.

In Kenya, several studies have indicated an emergence of the clonal spread of MDR KP and horizontal transfer of AMR genes (6,12–15). Increased use of 3rd generation cephalosporins in the early 2000s has increased extended-spectrum beta-lactamase (ESBL)-producing Enterobacterales, the most prevalent of which in Kenya are CTX-M-15 and SHV (10,12,16,17). With the rise in ESBL-producing Enterobacterales in Kenya, carbapenem use is increasingly leading to the emergence of carbapenem-resistant Enterobacteriaceae (CRE). In 2011, Poirel, et al. were the first to detect CRE in Kenya from urine samples bearing the New Delhi Metallo-β-lactamase (NDM-1) carbapenemase (15). In 2017, there were reports of KP isolates that encoded *bla_NDM-1_* and *bla_SPM_* carbapenemase genes (14) and isolates recovered from bacteremias bearing the pNDM-MAR-like plasmid backbone that has been shown to carry the *bla_NDM_* gene (6). More recently, Musila et al. (2021) identified KP isolates with the *bla_OXA-181_* carbapenemase gene. In addition, KP resistance to aminoglycosides, tetracyclines, chloramphenicol, and fluoroquinolones has also been observed in Kenya, typically associated with ESBL-production in MDR (6,10). Aminoglycosides are widely used to treat bacterial infections due to the ease of administration and access in Kenya (18). Musila et al. (2021) also reported the 16S rRNA methyltransferase genes, *rmtF*, and *rmtC*, in KP isolates which confer pan-aminoglycoside resistance. Resistance to chloramphenicol and sulfonamides is high in part due to its use as a first-line treatment option for enteric infections such as typhoid and HIV prophylaxis in Kenya (10).

In addition to antimicrobial resistance, KP possesses virulence genes that enhance its ability to cause infections, increase cell fitness, and evade the host immune system. KP colonizes host cells using adhesins such as fimbriae and pili (19). The production of a robust capsular polysaccharide confers resistance to host immune cells along with the O-antigen portion of the liposaccharide (LPS). KP attacks rival bacteria and eukaryotic cells by injecting potent endotoxins using type VI secretion machinery (20). Highly virulent KP increases the expression of *magA, rmpA*, and *rmpA2* genes linked to the mucoid phenotype for hypervirulence (21). Other mechanisms include the use of allantoin for a carbon and nitrogen source, the use of efflux pumps to eject antibiotics (22), and the capture of iron molecules from the host cells using siderophores (aerobactin, salmochelin, yersiniabactin) (23). Although some studies in Kenya have identified KP lineages ST11, ST15, and ST17, associated with hospital outbreaks (6,13), and the high-risk MDR ST147 (24), few studies have examined virulence genes in Kenyan KP isolates (13).

A clearer understanding of KP sequence types (STs) circulating within the community and healthcare systems across Kenya and the diversity and distribution of AMR and virulence genes would facilitate the monitoring and control of high-risk clones that are potentially hypervirulent and/or multidrug-resistant. Given the public health and clinical importance of KP and the knowledge gaps in Kenya, this study set out to examine KP isolates across Kenya and describe: 1) the sequence types and their distribution across Kenya, 2) the antimicrobial resistance phenotypes, and 3) the AMR and virulence genes present in the KP isolates.

## RESULTS

### Epidemiological and clinical characteristics of *K. pneumoniae* isolates

Clinical samples were collected from skin and soft tissue infections (SSTIs) (64%, 57/89) and urinary tract infections (UTIs) (29%, 26/89), and the hospital environment (7%, 6/89) from five counties in Kenya: Kisumu (39%, 35/89), Nairobi (26%, 23/89), Kisii (17%, 15/89), Kilifi (10%, 9/89), and Kericho (8%, 7/89). Sixty-two isolates were isolated from community-acquired (CA) infections, while 21 were from healthcare-associated (HA) infections. Thirty-seven isolates (42%) are MDR while 52 (58%) were non-MDR.

Of the four members of the *K. pneumoniae* Species Complex (2), *K. pneumoniae subsp. pneumoniae* represented the largest proportion of all isolates at 79% (70/89), with 44 recovered from SSTIs, 20 recovered from UTIs, and 6 from the hospital environment. This phylogroup had the largest number of MDR isolates (39%). The second most represented phylogroup was *K. variicola subsp. variicola* (18%, 17/89) with 15 isolates recovered from SSTIs and 2 isolated from UTIs. There was only one MDR isolate (kkp059) in this category. The least represented phylogroups were *K. quasipneumoniae subsp. quasipneumoniae* (2%, 2/89) and *K. quasipneumoniae subsp. similipneumoniae* (1%, 1/89) (Figure 1). Among the isolates from these minor subspecies, kkp022 and kkp034 were isolated from a UTI, while kkp036, the only MDR in this category, was recovered from an SSTI. All *K. quasipneumoniae subsp. quasipneumoniae, K. quasipneumoniae subsp. similipneumoniae, and K. variicola subsp. variicola*, except for one isolate (kkp078), were from community-acquired infections from different geographical locations (Figure1). There was no evident geographical clustering of all the phylogroups by county or infection types.

**Figure 1:**
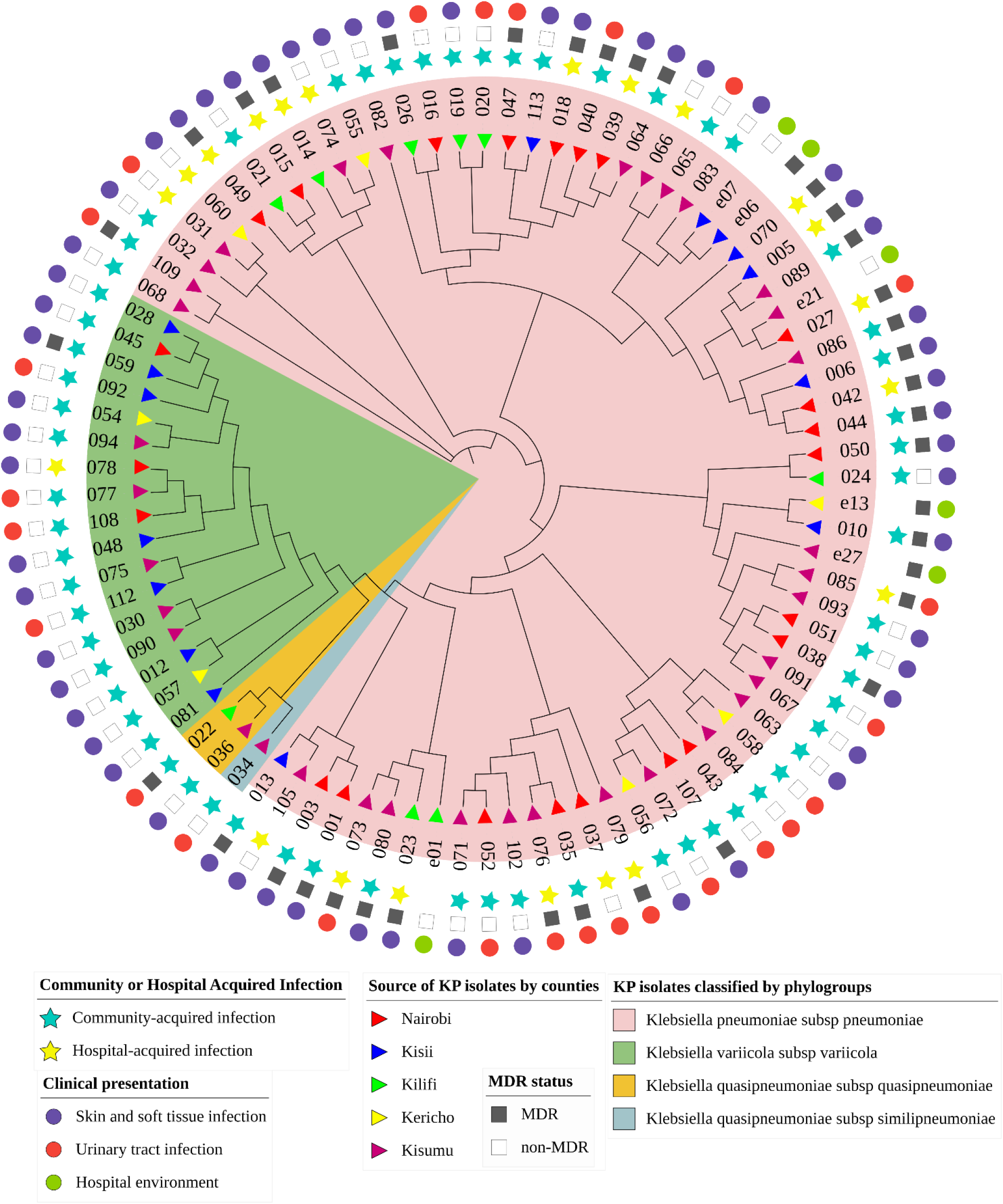
Circular cladogram showing epidemiological and clinical characteristics of *K. pneumoniae* isolates (n= 89). The color shading indicates the clustering of the isolates by phylogroups: *Klebsiella pneumoniae subsp. pneumoniae* (pink), *Klebsiella quasipneumoniae subsp. similipneumoniae* (blue), *Klebsiella quasipneumoniae subsp. quasipneumoniae* (gold), and *Klebsiella variicola subsp. variicola* (green). The triangle symbols in the innermost ring represent the geographic source of the isolates: Nairobi (red), Kisii (blue), Kilifi (green), Kericho (yellow), and Kisumu (purple). The star symbols represent the isolates recovered from community-acquired (blue) or healthcare-associated infections (yellow). The square symbols represent the multidrug resistance status of the isolates as either multidrug-resistant (black) or non-multidrug-resistant (white). The circle symbols in the outermost ring represent the clinical presentation of the isolates, i.e., skin and soft tissue infections (purple), urinary tract infection (red), and the hospital environment (green).

### Genomic characteristics of *K. pneumoniae* isolates

Draft genomes were generated from the 89 isolates: 40 via Illumina short-read sequencing and 49 via MinION-based long-read sequencing. The sizes of the draft genomes ranged from 5.2 to 5.9 Mb with an average G+C content of 57.27%, typical of *K. pneumoniae* genomes (38) (Table S1). The average N50 for the short and long reads was 226,402 and 5,147,285 base pairs, respectively. The isolates had 0 - 8 plasmid replicons, averaging 3 per isolate. The highest number of plasmid replicons were in genomes kkp001 (8), kkp018 (8), and kkp0e21 (8) (Table S2), whereas five genomes had no predicted replicons: kkp012, kkp030, kkp070, kkp102, and kkp112. The most abundant plasmids replicon types identified belonged to the Col and Inc family (particularly the IncF type) (Figure 2). The other Inc-like plasmid replicons identified were IncR, IncH, IncX, IncC, IncN, IncM, and IncY. Seven types of Col plasmid replicons, the second most represented type, were identified, dominated by Col(pHAD28) (Figure 2). Other plasmid types identified included pKP1433 and rep_KLEB_VIR in kkp056 in kkp043 genomes, respectively.

**Figure 2.**
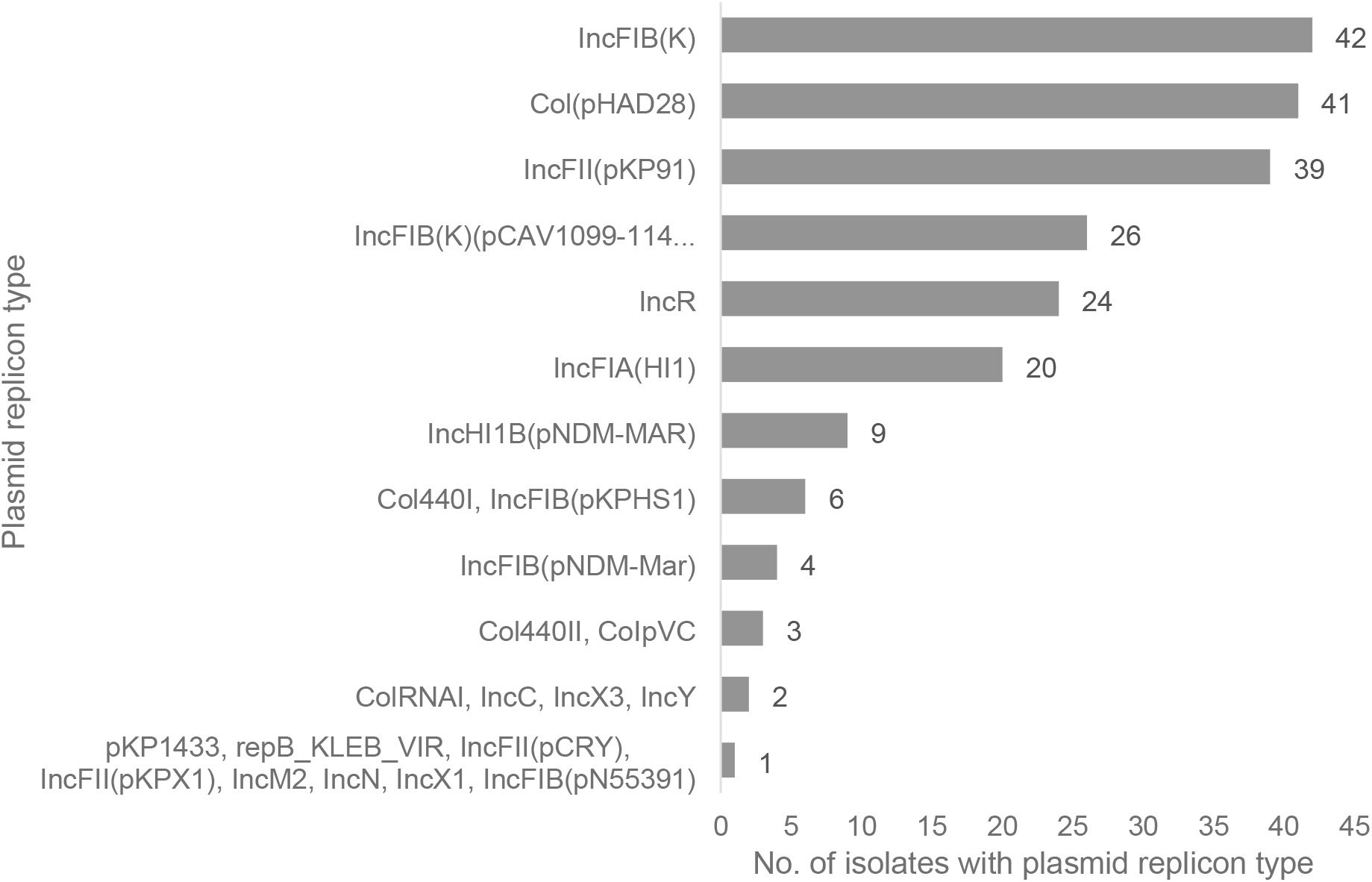
The types and number of plasmid replicons identified in the Kenyan *K. pneumoniae* isolates (N=89)

### Multilocus sequence types and distribution of *K. pneumoniae* isolates

The STs could only be assigned to 57% (51/89) of the isolates. The 38 unassigned isolates were not typeable due to assembling errors or inadequate genome coverage. The multilocus sequence types (STs) of the KP isolates were diverse, with thirty-eight different STs identified. STs that were represented by more than one isolate were ST15 (4/89), ST17 (3/89), and ST607 (3/89 each); and two isolates each represented the ST14, ST37, ST39, ST48, ST147, and ST307 lineages (Figure 3). The remaining 29 sequence types were represented by a single isolate and were geographically distributed as follows: Kisumu (n=17) - ST20, 45, 101, 198, 336, 391, 751, 1927, 2010, 3717, 3397, 3609, 5594, 5595, 5596, 5598, 5599; Nairobi (n=6) - ST55, 219, 966, 1786, 3692, 5600; Kisii (n=4) - ST25, 711, 3717, 5597 and Kilifi (n=2) - ST364 and 5593. Eight isolates had novel allelic profiles assigned and deposited in the *K. pneumoniae* MLST database (https://bigsdb.web.pasteur.fr/) (Table 1). There was no evident clustering of STs by infection type or ESBL and MDR status. Kisumu had the highest number of distinct STs (23), followed by Nairobi (12), Kisii (6), Kilifi (3), and Kericho (2). The high-risk lineages, ST14, ST15, ST17, ST307, and ST607, were concentrated in the biggest cities – Kisumu and Nairobi. Global high-risk strains ST14, ST15, ST307, and ST607, were predominantly from the *Klebsiella pneumoniae subsp. pneumoniae*.

**Figure 3.**
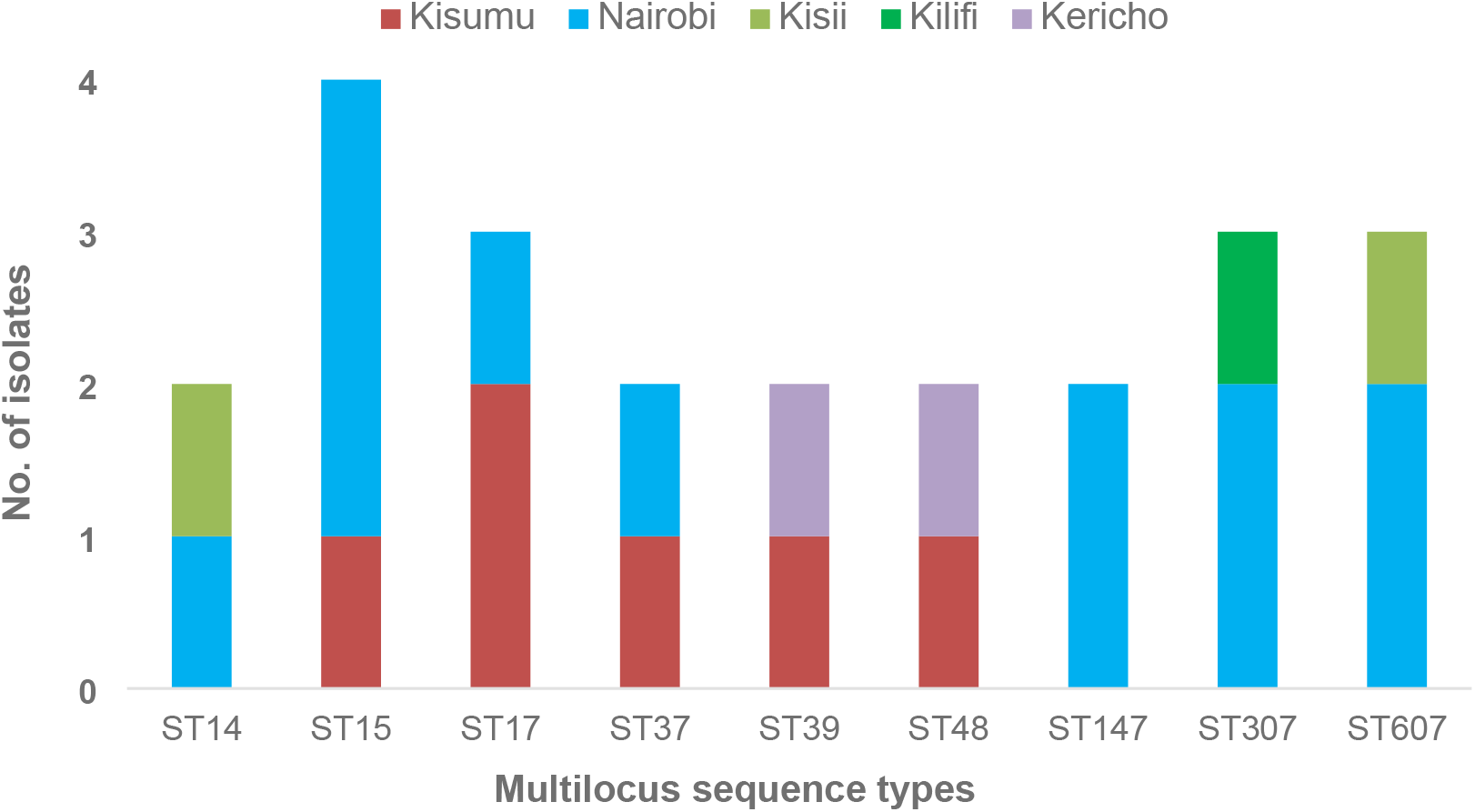
Distribution of the most abundant sequence types of the *K. pneumoniae* isolates by county.

**Table 1.**
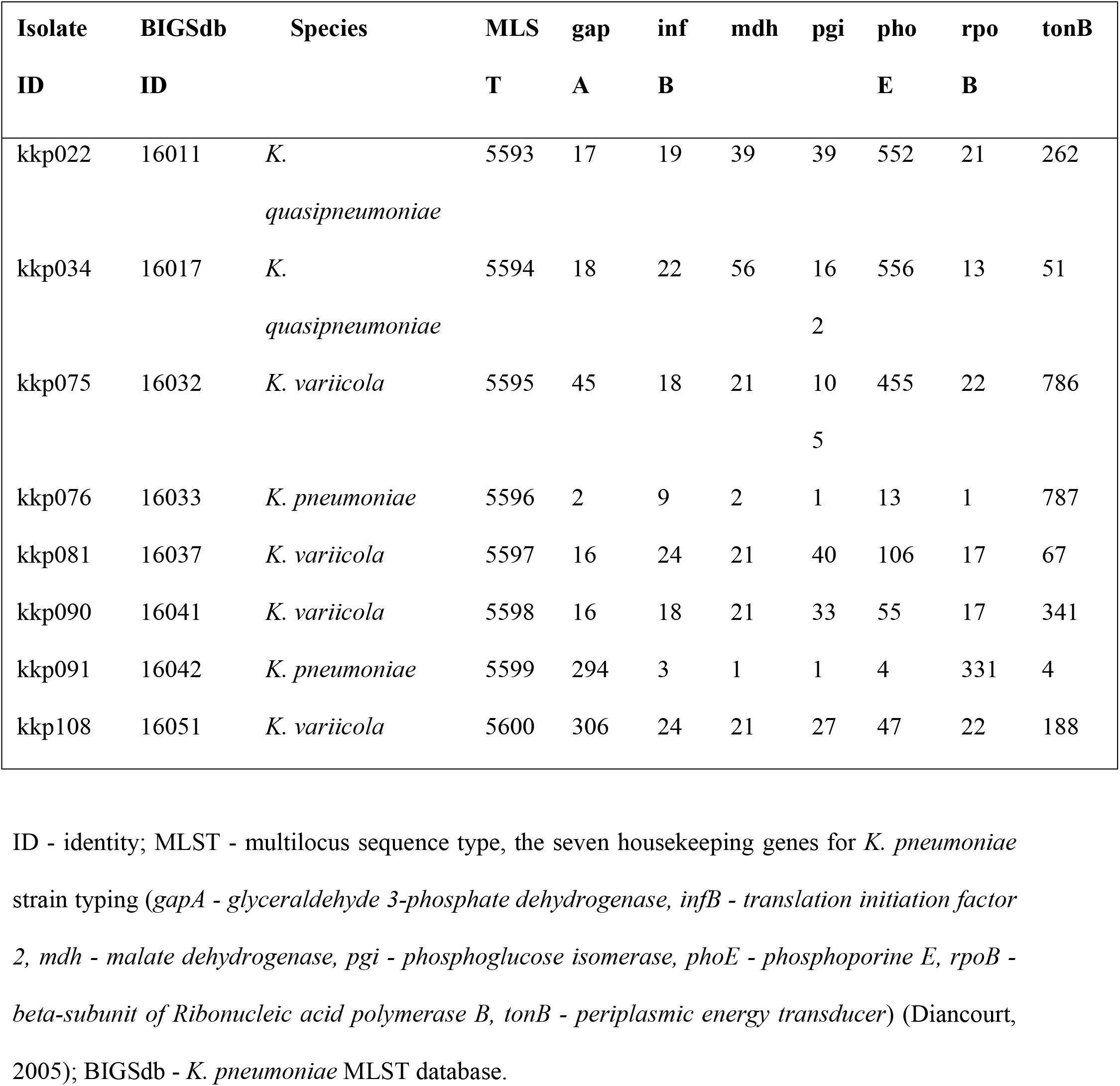
Allelic profiles of *K. pneumoniae* isolates with novel multilocus sequence types

### Antimicrobial Resistance

#### Phenotypic resistance profiles

Antibiotic susceptibility tests (AST) performed on the VITEK2^®^ platform (Table S3) identified high levels of non-susceptibility to trimethoprim (57%, 51/89), ticarcillin/clavulanate (49%, 44/89), cefuroxime axetil (49%, 44/89), cefuroxime (48%, 43/89), cefixime (48%, 43/89), ceftriaxone (47%, 42/89), cefepime (46%, 41/89), aztreonam (47%, 42/89), tetracycline (37%, 33/89) and minocycline (34%, 31/89); moderate non-susceptibility to levofloxacin (19%, 17/89), moxifloxacin (20%, 18/89) and chloramphenicol (19%, 17/89); and low levels of non-susceptibility to tigecycline (4%, 4/89) and meropenem (2%, 2/89). All isolates were non-susceptible to piperacillin (Figure 4). Among the dataset, 41% (37/89) were classified as MDR, i.e., resistant to 3 or more drug classes. A large proportion (47%, 42/89) were ESBL-producing isolates (Figure 4), of which 30 isolates were MDR. One isolate, kkp001, was pan-resistant (resistant to all 16 antibiotics tested in the panel).

**Figure 4.**
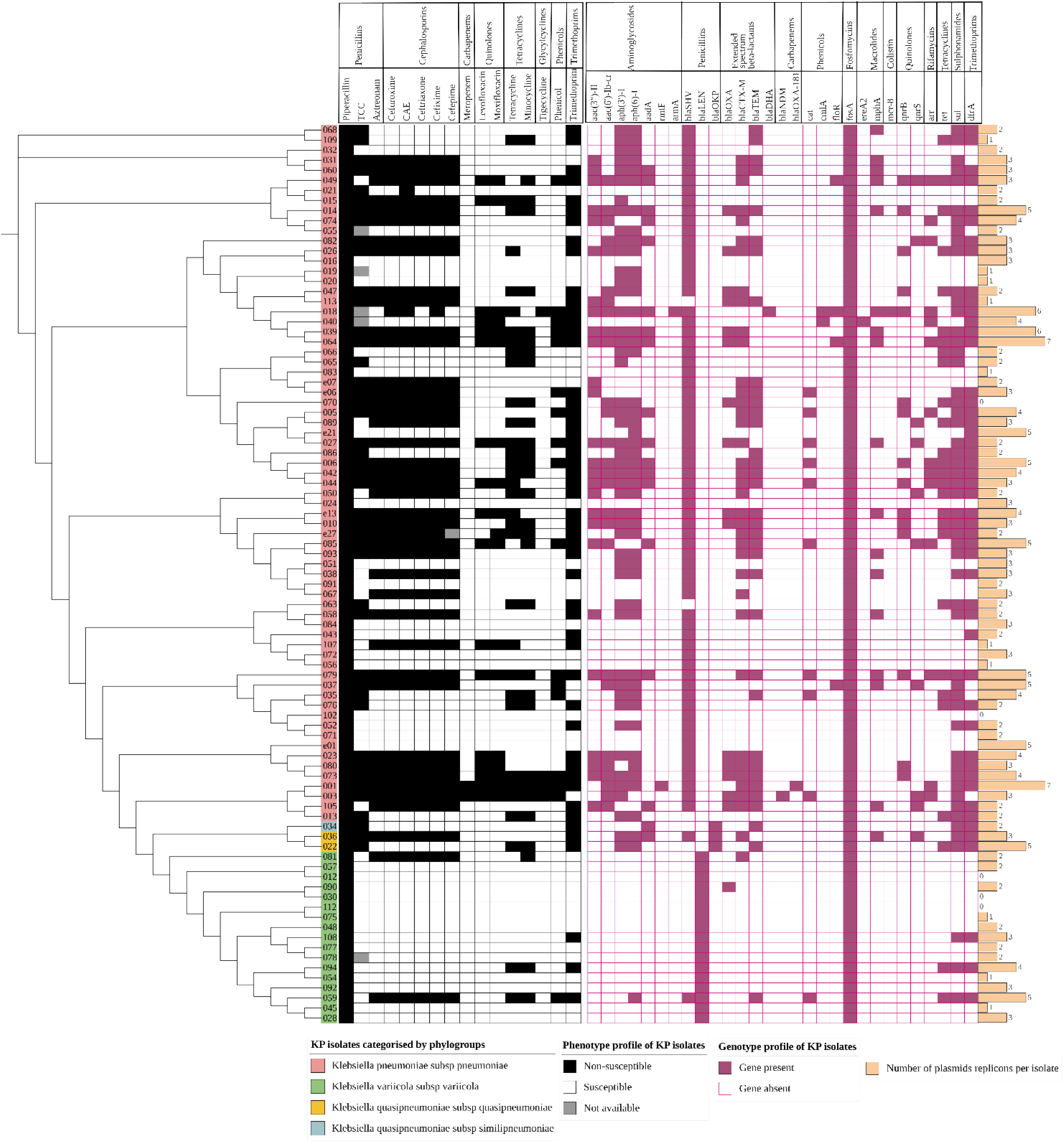
A heatmap of the phenotypic and genotypic profiles and plasmid replicon abundance of *K. pneumoniae* isolates (n=89). The cladogram on the left with colored labels on the edge indicates the clustering of the isolates by phylogroups: *Klebsiella pneumoniae subsp. pneumoniae* (pink), *Klebsiella quasipneumoniae subsp. similipneumoniae* (blue), *Klebsiella quasipneumoniae subsp. quasipneumoniae* (gold), and *Klebsiella variicola subsp. variicola* (green). The phenotypic profile is represented as an isolate being non-susceptible (black) or susceptible (white) to the antibiotic indicated on the bottom column header (and the drug class it belongs to on the top column header). The grey square indicates an isolate whose phenotypic result was not available. The genotypic profile is represented as a gene present (purple) or absent (white with purple outline). The genes are indicated on the column header and the drug class on the top column header. The beige bar plot represents the number of plasmid replicons found in the isolates. CAE – cefuroxime axetil. TCC – ticarcillin/clavulanate.

#### Genetic determinants of resistance

Comparison of the isolates’ phenotypic and genotypic antimicrobial susceptibility results demonstrated high concordance in the trimethoprim, beta-lactam, and tetracycline antibiotic classes, as demonstrated in Figure 4. In particular, there was strong concordance among the KP isolates between the presence of an ESBL-CTX-M gene and non-susceptibility to beta-lactams and cephalosporins (Figure 4), *dfrA* gene with trimethoprim non-susceptibility, and *tet* genes with tetracycline non-susceptibility. Most of the genomes generated from long reads enabled the detection of circularized plasmids bearing AMR genes (Table 2). MDR isolates had more plasmid replicons than the non-MDR isolates which had only 0 - 3 plasmid replicons (Figure 4).

**Table 2.**
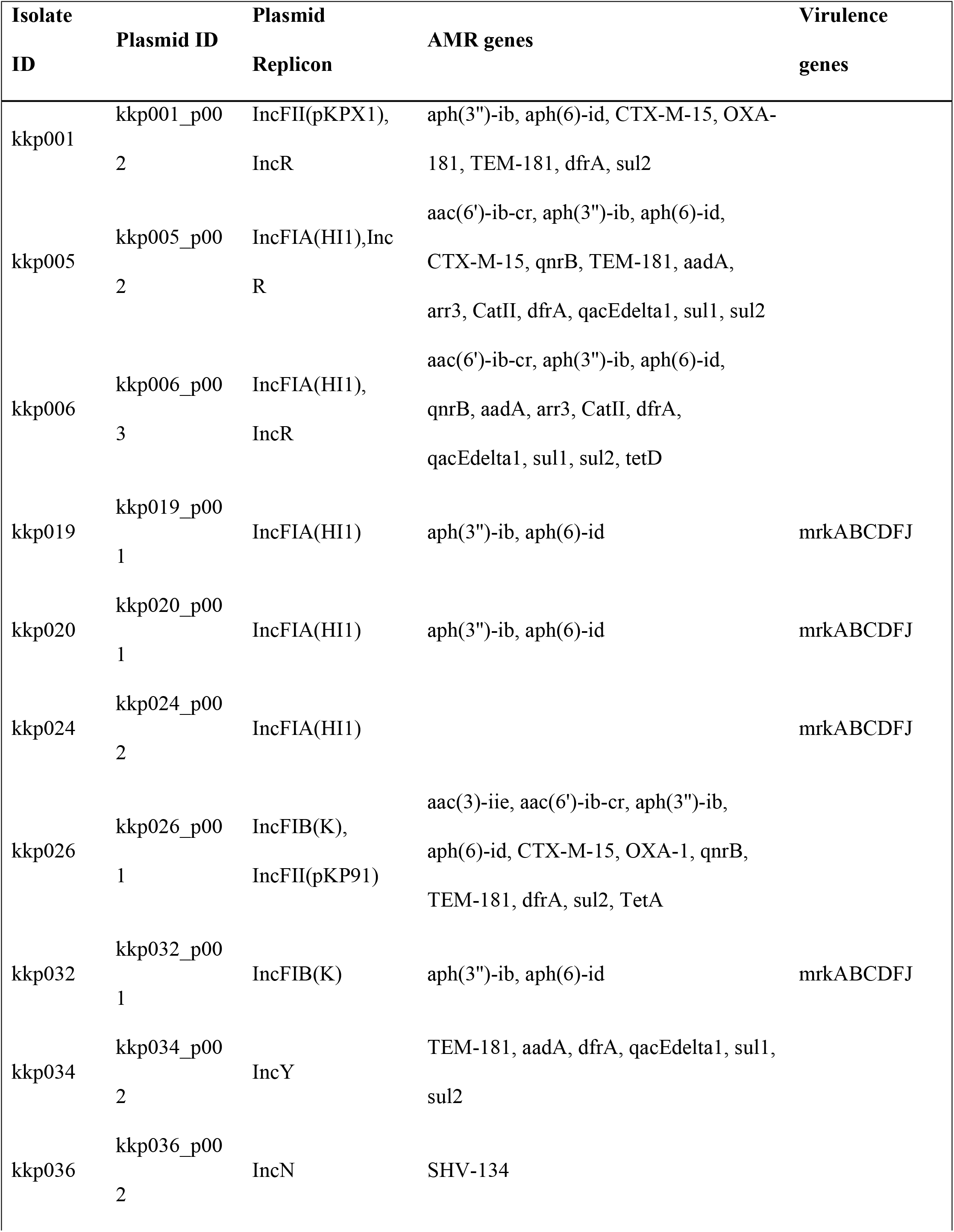
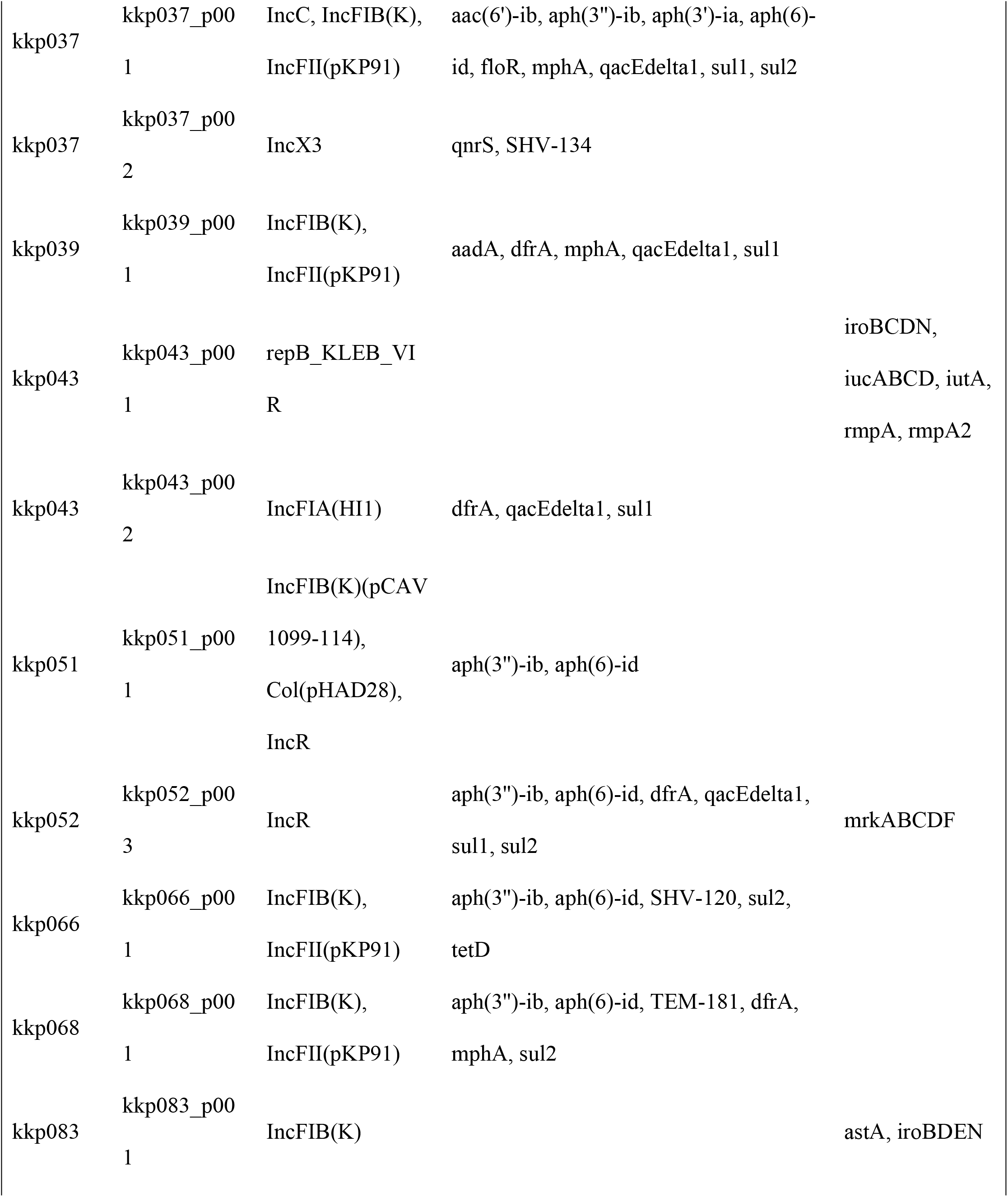
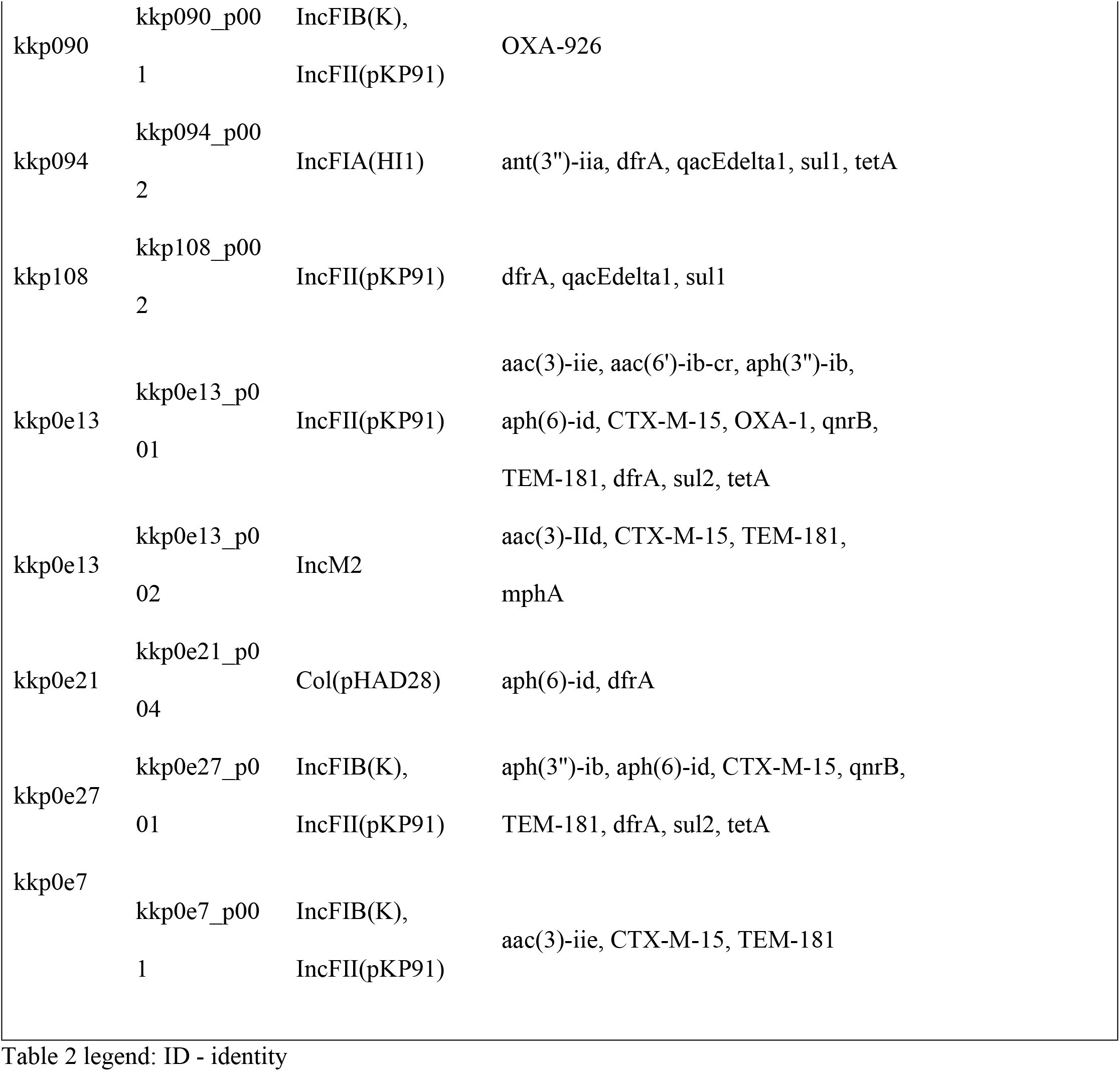
Antimicrobial resistance and virulence genes identified in the plasmids of the *K. pneumoniae* isolates

##### Beta-lactamase resistance genes

In the beta-lactam antibiotic class, genes for penicillinases, ESBLs, and carbapenemases were identified, summarized in Table S4. Each phylogroup demonstrated a distinct chromosomal penicillinase (Table S4), i.e., *Klebsiella pneumoniae* contained bla_SHV_, *Klebsiella quasipneumoniae* contained bla_OKP_, and *Klebsiella variicola* contained bla_LEN_, consistent with previous studies (8). However, some isolates carried a different additional penicillinase which was observed to have been acquired via a plasmid, e.g., kkp036 (*K. quasipneumoniae*) obtained a bla_SHV-134_ from an IncN plasmid (Table 2). There was a high diversity of plasmid-mediated ESBL genes in the isolates: bla_CTX-M-15_ (40%, 36/89), bla_CTX-M-98_ (1%, 1/89), bla_CTX-M-14_ (1%, 1/89), bla_TEM181_ (39%, 35/89), bla_OXA_ (18/89), and the less common bla_LAP2_ and bla_SCO1_. Two MDR isolates had carbapenemase genes: bla_NDM1_ (kkp003) and bla_OXA181_ (kkp001). Isolates kkp013 and kkp063 did not demonstrate any β-lactamase genes, and they were unsurprisingly ESBL-negative and non-MDR. Most isolates possessed at least one and at most six β-lactamase genes (Table S4). There were 42 ESBL-positive isolates, and 38 of them had a bla_CTX-M_ co-harbored with two or more other types of ESBL genes (Figure 4).

##### Non-beta-lactamase resistance genes

A previously described (11) colistin-resistant isolate, kkp018, carried an *mcr-8*, a bla_DHA1_, and 20 additional AMR genes (Table S4). Sixty-seven percent of the isolates (60/89) carried aminoglycoside-modifying enzymes (AME) genes i.e. *ant(3’*) (or *aadA*) encoding aminoglycoside nucleotidyltransferases; *aph(3’), aph(4*), and *aph(6*) encoding aminoglycoside phosphotransferases; *rmtF* and *armA* encoding16S rRNA methyltransferases; and *aac(3*) and *aac(6’)-Ib-cr* encoding aminoglycoside acetyltransferases (Figure 4, Table S4). The *aac(6’)-Ib-cr* gene also induces fluoroquinolone resistance. The genes associated with panaminoglycoside resistance were found in two isolates: *armA* (kkp018) and *rmtF* (kkp001). The genes for efflux pumps associated with aminoglycoside resistance (*arnT, crcB, acrD, baeR, cpxA*) were well-conserved among all the isolates. In addition,the common MDR efflux pump genes were identified: *LptD, CRP, H-NS, KpnEFGH, acrAB, marA, mdtBC, msbA*, and *ramA*.

Three mechanisms for quinolone resistance were detected. First, plasmid-mediated quinolone resistance (PMQR) genes (*qnrB* or *qnrS1*) identified in 26 isolates. Second, chromosomal-encoded efflux pumps (*emrR, oqxA*, and *oqxB*) genes constitutive in all isolates. Third, gene mutations were detected in gyrase A (*gyrA*) - Ser83Phe, Asp87Aspn, and Ser83Ile - and Deoxyribonucleic acid topoisomerase IV subunit A (*parC*) - Ser80Ile (Table S6), which are involved in DNA synthesis. Twenty-nine isolates (32%, 28/89) had the tetracycline-resistance genes, *tetA* or *tetD*. Fifteen isolates (17%, 15/89) carried the chloramphenicol resistance genes *cat1, catII*, and *catB3* (encoding chloramphenicol acetyltransferases) as well as *floR* and *cmlA* (encoding chloramphenicol efflux pumps). Resistance to sulphonamide and trimethoprim, administered as co-trimoxazole, was mediated by the *dfrA* trimethoprim resistance gene and *sul* sulphonamide resistant gene, found in 54 and 55 isolates, respectively. Resistance to other drug classes was conferred by: fosfomycins - *fosA*; macrolides - *mphA, mphE, msrE*, and *ereA2*; rifamycins - *arr2* and *arr3*; and even antiseptics - *qacEΔ1* and *qacL*.

#### Virulence factors associated with the *K. pneumoniae* isolates

Known virulence factors involved in adherence, biofilm formation, capsule synthesis regulation, mucoid phenotype regulation, immune evasion, secretion system, serum resistance, siderophores expression (enterobactin, yersiniabactin, aerobactin, and salmochelin), efflux pump expression, allantoin utilization, and enterotoxin generation were detected among the isolates (Figure 5). The most ubiquitous were the chromosomal genes *fim, mrk*, and *ecp* for adherence and biofilm formation, which were present in all isolates except kkp043 which lacked the *fim* genes. In addition, some isolates carried multiple copies of the *mrk* genes in plasmids (Table 2). Other genes identified in all the isolates were those for serum resistance factors that determine the ‘O-antigen’ lipopolysaccharide serotype, the immune evasion factors which determine the polysaccharide capsule (*K antigen*) type, capsule synthesis regulation (*rcs*), efflux pump expression (*acrAB*), and enterobactin (*ent, fep*) (Table S5). In addition, all isolates possessed type VI secretion system loci genes except kkp034.

**Figure 5.**
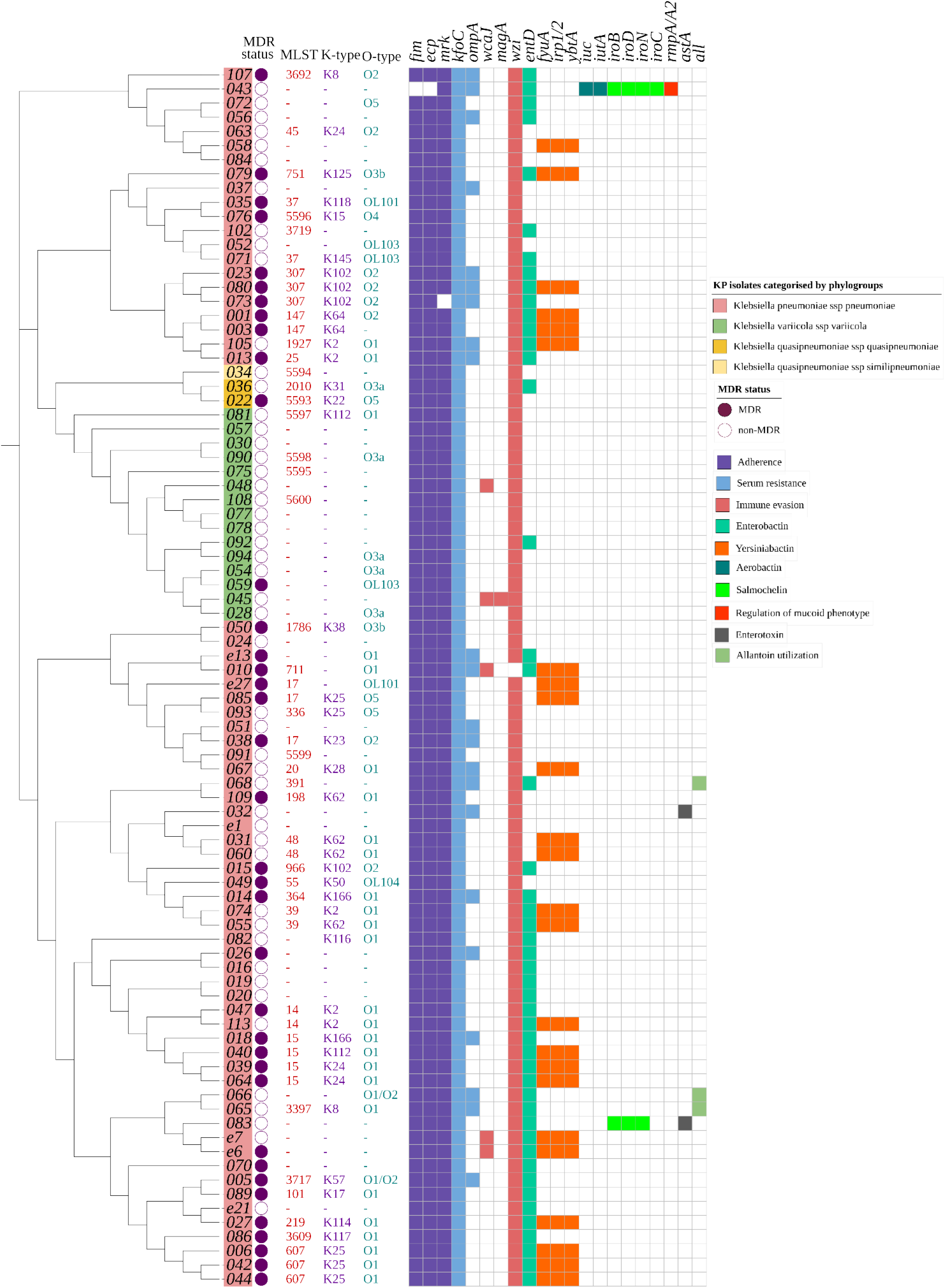
A heatmap of the multidrug resistance status, ST, serotype, and virulence gene content of *K. pneumoniae* isolates (n=89). The cladogram on the left with colored labels on the edge indicates the clustering of the isolates by phylogroups: *Klebsiella pneumoniae subsp. pneumoniae* (pink), *Klebsiella quasipneumoniae subsp. similipneumoniae* (blue), *Klebsiella quasipneumoniae subsp. quasipneumoniae* (gold), and *Klebsiella variicola subsp. variicola* (green). The circular symbols represent the multidrug resistance status of the isolates as either multidrug-resistant (purple) or non-multidrug-resistant (white). The serotypes assigned to the isolates are indicated as follows: multilocus sequence type (MLST) (red), capsule type (purple), and O type (green). The virulence profile is represented as a gene present (color) or absent (white) for factors: adherence (purple), serum resistance (blue), immune evasion (pink), enterobactin (green), yersiniabactin (orange), aerobactin (teal), salmochelin (luminous green), regulation of mucoid phenotype (red), enterotoxin (gray), and allantoin utilization (pale green). (-) indicates unassigned serotypes.

This study identified no evident clustering of capsule and lipopolysaccharide types based on geographical locations or clinical presentation. There were 10 different O-loci types identified (O1, O2, O3a, O3b, O4, O5, OL101, OL103, and OL104) in the 57 assigned isolates, whereby O1 (46%, 26/57) and O2 (19%, 11/57) were the most common and clinically significant. There were 24 different K-types in the 45 assigned isolates, and the most abundant were K2 (11%, 5/45), K25 (11%, 5/45), K102 (9%, 4/45), K62 (9%, 4/45), and K24 (7%, 3/45).

The ybt loci, identified in 24/89 isolates, encode for yersiniabactin siderophores found within conjugative transposons in the chromosome. Based on a new typing scheme (39) updated in the kleborate pipeline (34), this study identified four distinct *ybt* types: *ybt14* found within ICEKp5, *ybt15* in an ICEKp11, *ybt16* in an ICEKp12, and *ybt9* within an ICEKp3. In addition, the analysis revealed three isolates (Figure 5) with chromosomally encoded genes for allantoin utilization and two isolates (kkp012 and kkp045) (Figure 5) with *magA* and *K2* capsule types linked to hypervirulence (40).

Several isolates were unique in having plasmid-encoded virulence genes. For example, a hypervirulent isolate (hvKP), kkp043, was identified bearing the repB_KLEB_VIR plasmid containing *rmpA* and *rmpA2* genes, which regulate the expression of the mucoid phenotype, salmochelin (*iroBCDN*), and aerobactin (*iucABCD, iutA*). In addition, Kkp083 also demonstrated an *iroBDEN* cluster and a heat-stable enterotoxin (*astA*) gene carried in an IncF(K)_1 plasmid, in contrast to the chromosomally-bound *astA* gene in kkp032. This study did not identify MDR hypervirulent isolates with AMR and hypervirulent genes (41). Furthermore, the MDR isolates did not carry factors associated with hypervirulence, while the hypervirulent isolates were mostly antibiotic susceptible, i.e., they were non-MDR and ESBL-positive (Figure 5).

## DISCUSSION

This study characterized 89 isolates based on their clinical, geographic, genotypic, and phenotypic characteristics. It was noted that all the four recognized phylogroups, were represented with *Klebsiella pneumoniae subsp. pneumoniae* isolates predominating, consistent with findings from other studies (42–44). Some differences were observed in the characteristics of the phylogroups. For example, *Klebsiella variicola subsp. variicola* isolates are typically linked to bloodstream infections and UTIs (45,46); however, those identified in this study were mainly associated with SSTIs (Figure 1) and were largely antibiotic susceptible (Figure 4). In previous studies, *Klebsiella quasipneumoniae subsp. similipneumoniae* isolates were linked to nosocomial infections such as UTIs (47,48). Yet, in this study, one isolate, kkp034, caused a community-acquired SSTI, potentially indicating a broader distribution of this phylogroup in Kenya as found in other countries, such as sewage in Brazil (49) and a turtle in China (50).

The lineages identified in this study were highly diverse, and most of them were local strains that have not been described in other countries. Globally disseminated high-risk lineages such as ST14, ST15, ST307, and ST607, were identified (Figure 3). These high-risk lineages are bacterial pathogens that easily acquire and disseminate antimicrobial resistance (51). For example, ST14, ST15, and ST147 have been linked to the spread of carbapenemase resistance genes in many countries (52,53). In this study, the two extensively drug resistant (XDR) ST147 strains carried a bla_OXA-181_ (kkp001) and a bla_NDM_ gene (kkp003), respectively, while the MDR ST14/15 strains carried several ESBL genes. Notably, kkp018 (ST15) harbored a mobile colistin-resistant gene, mcr-8, and an AmpC betalactamase gene, blaDHA (Figure 4). ST307 and ST607 are emerging strains linked to ESBL infections (Long, et al., 2017), and they were observed in six MDR strains isolated from SSTIs and UTIs (Figure 1 and 5). ST17 is a regional strain that has been implicated in outbreaks in Kilifi (6), Mwanza (54), and Kilimanjaro (55). This study identified three MDR and ESBL positive ST17 strains, isolated from a community-acquired SSTI, a nosocomial UTI, and a hospital environmental swab (Figures 1 and 5). High-risk clones have also been linked to specific serotypes, e.g., ST607-K25, responsible for a nosocomial outbreak at a neonatal intensive care unit in a hospital in France (56). Significantly, this study identified three MDR and ESBL-positive ST607-K25 clones associated with SSTIs (Figures 5).

The presence of these high-risk strains in Kenya indicate the significant clinical and public health threat they pose. However we noticed that this threat was highest in Kisumu and Nairobi, the two largest cities in the country, were most of the global lineages (79%, 11/14) were identified. These cities are travel hubs with large referral hospitals serving patients from a wide geographical area locally and global travelers. The diverse patient population could explain the concentration of local, regional and global lineages, the great strain diversity, and novel alleles in isolates from the two counties compared to Kisii, Kilifi, and Kericho counties.

Plasmids are the main vehicle for AMR gene transmission and in this study we found that a majority of the AMR genes were carried in plasmids, particularly IncFIB(K) and IncFII(pKP91) (Table 2), which are common among Enterobacterales. The predominant plasmid replicon types belonged to the diverse Incompatibility (Inc) family (57), whose host range is mainly limited to Enterobacterales (58) known to have large multi-replicon plasmids (59). The multi-replicon IncF plasmids contain the FII, FIA, and/or FIB replicons (Table 2 and S2) which account for their high abundance (Figure 2). Unsurprisingly, antibiotic susceptible KP isolates (kkp012, kkp030, kkp102, and kkp0112) contained few or no plasmids and were ESBL negative and non-MDR. The MDR kkp070 was exceptional because although it possessed no plasmid replicons, it had several resistance genes integrated into a genetic island in the chromosome. These integration events were not uncommon, as plasmid-associated AMR genes were detected in the chromosome of several isolates: kkp005 and kkp006 carried a bla_CTX-M-15_ gene, kkp109 had a dfrA14 trimethoprim resistant gene, and kkp001 possessed several AMR genes (*aac(6’)-Ib9*, bla_CTX-M-15_, *arr-2, and rmtF*). The integration of AMR genes into the chromosome is alarming because the resistance is transferred clonally, becomes part of the core genome, and increases the spread and prevalence of non-susceptible KP lineages.

Although the col plasmid family were the second most dominant group, the majority of col-like plasmids did not contain any resistance or virulence genes, except for the abundant colE1 type, Col (pHAD28), which typically carries qnrS1 and other AMR genes (58) as observed in kkp0e21 (Table 2). In addition, col plasmids benefit KP because they produce bacteriocins lethal to rival bacteria. One of the other plasmid families identified was the pK1433 (kkp056) and is associated with the blaKPC2 gene (60), which provides a mechanism for carrying and spreading KPC-type genes. The isolates with plasmids carrying a bla_CTX-M_ gene also carried bla_OXA_, bla_SHV_, or bla_TEM_ genes, implying that these resistance genes are transferred together (e.g., ESBL-positive kkp081 and kkp059 isolates possessed bla_CTX-M-15_, bla_SHV-134_, and bla_TEM181_ genes). The most intriguing isolates were the ESBL-positive isolates (kkp015, kkp070, and kkp107) which only contained the bla_SHV_ penicillinase gene. We hypothesize that the ESBL phenotypes in these isolates may have been due to alternative resistance mechanisms such as up-regulation of MDR efflux pumps. The expression of genes encoding MDR efflux pumps is noteworthy because their activity induces resistance of different antibiotic classes non-specifically and can cause phenotypic and genotypic discordance.

Examination of the virulence factors among the KP phylogroups highlighted one main difference. *K. quasipneumoniae subsp similipneumoniae* did not demonstrate the Type VI protein secretion system found in the other phylogroups (Table S5); instead, it possessed the Type II secretion as previously noted (61). Siderophores were the most significant virulence genes detected. The ubiquitous enterobactin scavenges iron from host cells; however, their activity is neutralized by human lipocalin-2 protein (62). In response, KP uses the more virulent yersiniabactin to bind iron and other heavy metals like copper to avoid metal toxicity and phagocytosis using reactive oxygen species (63). According to Holt, et al. (2015), acquiring yersiniabactin is usually the first step in accumulating more potent siderophores to make them more invasive. Yersiniabactin genes were found among the *K. pneumoniae subsp pneumoniae* isolates causing various clinical infections and from diverse geographical locations. Most of them were MDR and ESBL positive, making their infections persistent and antibiotic-resistant and likely contributing to their dominance over the other phylogroups.

Classical hypervirulent KP (hvKP) isolates are generally antibiotic susceptible (64), and they are characterized by having *rmpA/A2* and *magA* virulence factors as well as the *K2* capsule type (53). This study identified one potentially hypervirulent isolate (kkp043), which carried *rmpA* and *rmpA2* genes. In addition, it also carried genes for aerobactin and salmochelin that were carried on a repB_KLEB_VIR plasmid (Table 2). A trade-off between virulence and antibiotic resistance was demonstrated in this potentially hypervirulent isolate (kkp043) as it was non-MDR and ESBL-negative and had only one other plasmid bearing the IncFIA(HI1) replicon, *dfrA5*, and *sul1* genes (Table 2). Additional examples were the two isolates (kkp012 and kkp045) with *magA* genes which were ESBL-negative and non-MDR. Furthermore, non-MDR K2 strains possessed the ybt operon, which the MDR K2 strains lacked (Figure 5). Despite this, there is growing concern about the rise of MDR hvKP globally (39,65–68), which cause severe infections with few treatment options (69) emphasizing the significance of monitoring virulence characteristics of KP in Kenya.

The study had several limitations. The first was that the two sequencing technologies utilized had different strengths and weaknesses, which could introduce bias in the genomic analysis. The draft genomes from the long reads had fewer contigs (<=12) (Table S1) and were more contiguous, but they had less depth/coverage, while the short reads produced less contiguous draft genomes with greater depth (>40) (Table S1). More contiguity enabled better plasmids reconstruction, while sufficient depth enabled better assignment of multilocus sequence types. The second limitation was that susceptibility tests were carried out on the Vitek2^®^ platform using a limited number of antibiotics and were not verified using another phenotypic method. Nevertheless, there was good concordance between the phenotypic and genotypic results. Finally, *in vivo* tests were not conducted to confirm the virulence gene activity, so the virulence results are only predicted.

## CONCLUSION

The findings of this study contribute in several ways to our understanding of the genotypic and phenotypic characteristics of Kenyan *K. pneumoniae* isolates. The multi-center approach provided more nationally relevant data, unlike prior studies with fewer isolates from single sites. These findings describe a KP population with diverse sequence types, highly abundant and diverse resistance, and virulence profiles. In addition, the presence of high-risk clones in the major cities Nairobi and Kisumu enhances their transmissibility within and outside the country. Further research should be conducted to correlate the genotypic findings of virulence with phenotypic data, and additional analysis should investigate the genotypic environment of the acquired antimicrobial resistance genes to determine their risk of spread.

## MATERIALS AND METHODS

### Ethics Statement

Ethics approval for the study was obtained from the KEMRI Scientific and Ethics Review Unit, Nairobi, Kenya (KEMRI SSC# 2767) and the Walter Reed Army Institute of Research Institutional Review Board, Silver Spring, MD, USA (#2089). Written informed consent was obtained from each participant. The investigators have adhered to the policies for the protection of human subjects as prescribed in AR 70-25.

### Study Site

The bacterial culture, DNA extraction, quantification, and Oxford Nanopore sequencing were conducted in the Kenya Medical Research Institute (KEMRI) - Center for Microbiology Laboratory. Illumina sequencing was performed at the Multidrug-Resistant Organism Repository and Surveillance Network (MRSN), Walter Reed Army Institute of Research (WRAIR). The bioinformatic analysis was conducted using the high-performance computing server at the Department of Emerging Infectious Diseases, USAMRD-Africa in KEMRI, Nairobi.

### Study Samples

The study analyzed hospital environmental and clinical isolates of KP that were collected between May 2015 to March 2020 from eight hospitals in five counties in Kenya as part of an ongoing antimicrobial resistance surveillance study (KEMRI2767/WRAIR 2089) and an environmental study (KEMRI 3482/WRAIR 2416). The clinical samples were urine, wound swabs, and pus collected from consenting patients with suspected bacterial infections. The environmental samples were collected via swabs of high-touch areas in the participating hospitals. *Klebsiella pneumoniae* identification and antimicrobial susceptibility testing (AST) were performed on the Vitek2^®^ system (bioMérieux, France) using the GN-ID and XN05-AST cards. The AST panel consisted of penicillins (piperacillin and ticarcillin/clavulanic acid), monobactam (aztreonam), cephalosporins (cefuroxime, cefuroxime axetil, cefixime, ceftriaxone, and cefepime), carbapenems (meropenem), fluoroquinolones (levofloxacin and moxifloxacin), tetracyclines (tetracycline and minocycline), glycylcycline (tigecycline), phenicol (chloramphenicol) and trimethoprim. The AST results were interpreted according to CLSI guidelines (2018), and isolates were classified as either multidrug-resistant (resistant to 3 or more drug classes) or non-multidrug resistant and ESBL positive or negative.

### DNA Extraction and Sequencing

Forty isolates were sequenced on an Illumina MiSeq platform at MRSN-WRAIR as previously described (24). The remaining 49 samples were sequenced using the MinION platform (Oxford Nanopore Technology). Forty-nine KP isolates were inoculated on Mueller Hinton Agar plates and incubated for 24 hours at 37°C. *Klebsiella pneumoniae* JH930422.1 was included as a positive control. Approximately 14 × 10^8 cells/mL at OD 600 of overnight bacterial cells were resuspended in nuclease-free water and centrifuged at 5,400g for 10 minutes to pellet the cells. Total DNA was then extracted using the DNeasy® UltraClean Microbial Kit (QIAGEN Inc., Netherlands) according to the manufacturer’s instructions. DNA purity was determined on the Nanodrop One (ThermoFisher Scientific) and quantified on the Qubit dsDNA fluorometer. For library preparation, the extracted DNA was end-repaired using the NEBNext® UltraII End Repair/dA-Tailing kit using the manufacturer’s instructions. LSK 109 ligation kit, EXP-NBD 104 (1–12), and EXP-NBD 114 (13–24) Native Barcoding kits were used to barcode each DNA sample and adapters added. The DNA library was loaded onto a FLO-MIN106 R9.4.1 flowcell for sequencing based on the standard Oxford Nanopore Technology (ONT) 1D sequencing protocol launched in the MinKNOW v20.10.3 software. A quality control experiment was conducted to evaluate the Nanopore workflow by sequencing the lambda phage DNA on the FLO-MIN106 R9.4.1 flowcell for 6 hours as per the Lambda DNA control experiment protocol, and the sequences compared with the reference sequence (NC_001416.1). Guppy v4.4.2 was used to basecall and trim the barcodes and adapters from both ends of the reads.

### De novo assembly of raw reads and database querying

The fastQ files were filtered to retain only those with a Q-score ≥7. The previously sequenced paired-end Illumina reads were retrieved from the in-house server for downstream analysis. The adapters in the short pair-ended reads were trimmed using *Trimmomatic v0.39* (25). The trimmed Illumina reads were assessed for quality using *FastQC v0.11.9* (26) before de-novo assembly using *the default Shovill v1.1.0* (27) *pipeline* settings. Next, the draft assemblies were polished using *pilon v1.24* (28). Finally, the ONT long-reads were de-novo assembled using *flye* assembler v2.8.1 (29) with the plasmid option, followed by one polishing round using *medaka* v1.3.2. All polished draft assemblies from Illumina and ONT sequencing were analyzed in the same way. First, the quality was assessed using *QUAST*(30). Then, the draft assemblies were queried using the *ABRicate* pipeline v1.0.1 against *CARD* (31) to identify AMR genes, *VFDB* (32) to identify virulence factors, and *PlasmidFinder* (33) to identify plasmids. The assemblies were queried against the *Klebsiella MLST* database using the command line *mlst v2.19* pipeline to determine the sequence types, while the capsule (K) and O types were determined using the *Kleborate* pipeline (34) against the *Kaptive* database (35). Next, a new ybt typing scheme updated in the *Kleborate* pipeline was explored that assigned allelic profiles to yersiniabactin genes (34). Finally, a maximum-likelihood phylogenetic tree was generated using *Parsnp v1.2* (36) and NC_009648.1 as the reference genome. The tables were created using *flextable* (R package), while the circular tree and heatmaps were generated using Interactive Tree of Life (*iToL) v6.3.2* tree annotator (37).

## DECLARATIONS

### Data summary

All data generated or analyzed during this study are included in this published article and in the supplementary data files. Six supplementary tables are available on the online Supplementary Material of this article. Eighty-nine genome assemblies are deposited in the NCBI GenBank database under BioProject PRJNA777842. Novel allelic profiles of eight isolates are assigned and deposited in the *K. pneumoniae* MLST database (https://bigsdb.web.pasteur.fr/)

## Acknowledgments

The authors would like to acknowledge the Kenya Medical Research Institute and the United States Army Medical Research Directorate -Africa staff who carried out the patient sampling, biochemical and antimicrobial susceptibility tests, and the study participants at the participating hospitals. The authors also acknowledge the Walter Reed Army Institute of Research-Multidrug-Resistant Organism Repository and Surveillance Network staff who performed the Illumina sequencing and the Institut Pasteur teams for curating and maintaining the BIGSdb-Pasteur databases at http://bigsdb.pasteur.fr/.

## Conflicts of Interests

All authors declare they have no competing interests or conflicts of interest.

## Funding information

This work was funded by the Armed Forces Health Surveillance Division, Global Emerging Infections Surveillance Branch (PROMIS ID 20160270153 FY17-20). The study funders had no role in the study design, data collection, analysis, decision to publish, or preparation of the manuscript.

## Disclaimer

The material for this publication has been reviewed by the Walter Reed Army Institute of Research, and there is no objection to its publication. The opinions or assertions contained herein are the private views of the authors and are not to be construed as official or as reflecting the views of the Department of the Army or the Department of Defense. The investigators have adhered to the policies for the protection of human subjects as prescribed in AR 70-25.

## Author Contributions

A. W. M – conceptualization, methodology, formal analysis, investigation, writing – original draft preparation, visualization. Cecilia K. – conceptualization, methodology, visualization. S. K. – methodology, visualization. H. J. S. – writing – original draft preparation. Caleb K. – supervision. M. J. M. – methodology. J. K. – supervision. L. M. – conceptualization, resources, writing – original draft preparation, supervision, funding.

## LIST OF ABBREVIATIONS

AME: aminoglycoside modifying enzyme
AMR: antimicrobial resistance
AST: antimicrobial susceptibility test
CAI: community-acquired infection
CARD: Comprehensive Antibiotic Resistance Database
CRE: carbapenemase resistant enterobacteriaceae
ESBL: extended spectrum beta lactamase
HAI: health-acquired infection
KEMRI: Kenya Medical Research Institute
KP: *Klebsiella pneumoniae*
LPS: lipopolysaccharide
MDR: multidrug resistance
MLST: multilocus sequence type
MRSN: Multidrug-Resistant Organism Repository and Surveillance Network
NCBI: National Center for Biotechnology Information
NDM: New Delhi metallo beta lactamase
ONT: Oxford Nanopore Technology
PMQR: plasmid-mediated quinolone resistance
SSTI: skin and soft tissue infection
ST: sequence type
UTI: urinary tract infection
WHO: World Health Organisation
WRAIR: Walter Reed Army Institute of Research
XDR: extensively drug-resistant

## Supplementary Material captions

Supplementary Table 1 - Genomic and epidemiological characteristics of *K. pneumoniae* isolates

Supplementary Table 2 - Plasmids replicons present in the *K. pneumoniae* isolates

Supplementary Table 3 - Phenotypic antimicrobial susceptibility test (AST) results of the *K*.

*pneumoniae* isolates

Supplementary Table 4 - Antimicrobial resistance genes in the *K. pneumoniae* isolates

Supplementary Table 5 - Virulence genes identified in the *K. pneumoniae* isolates

Supplementary Table 6 - Fluoroquinolone mutations identified in the *K. pneumoniae* isolates

---

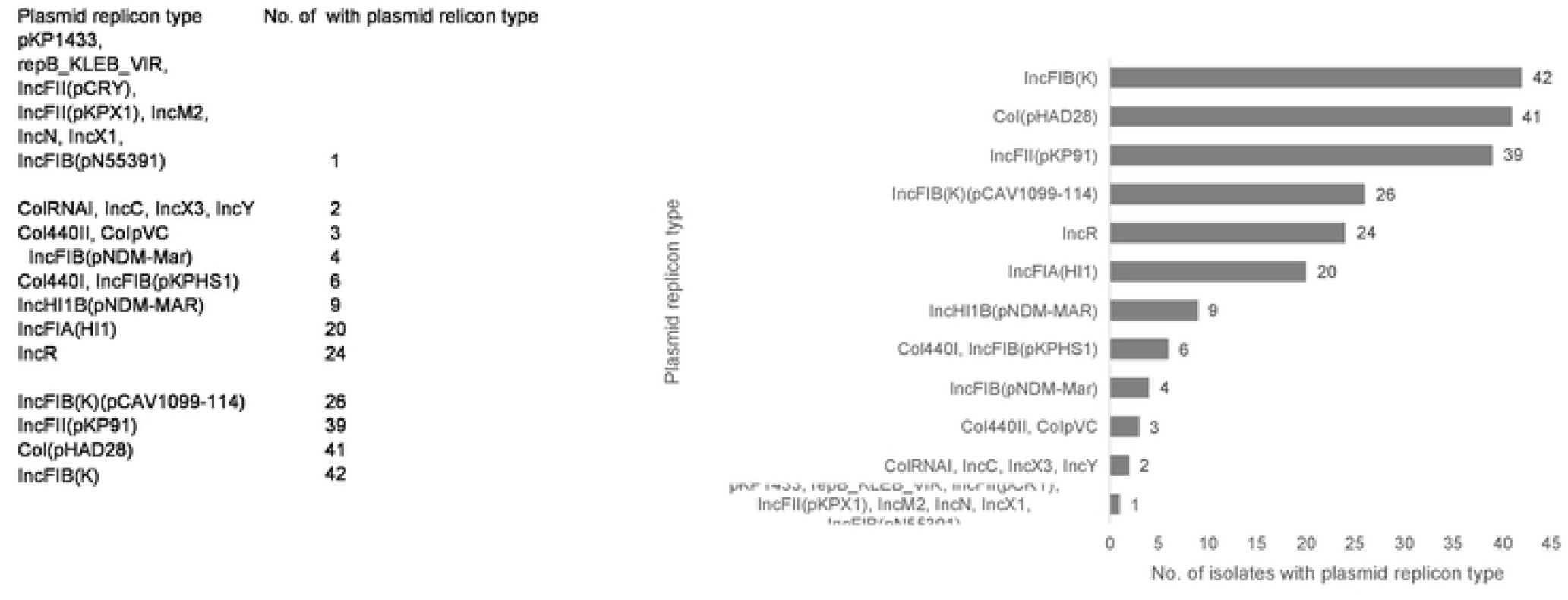

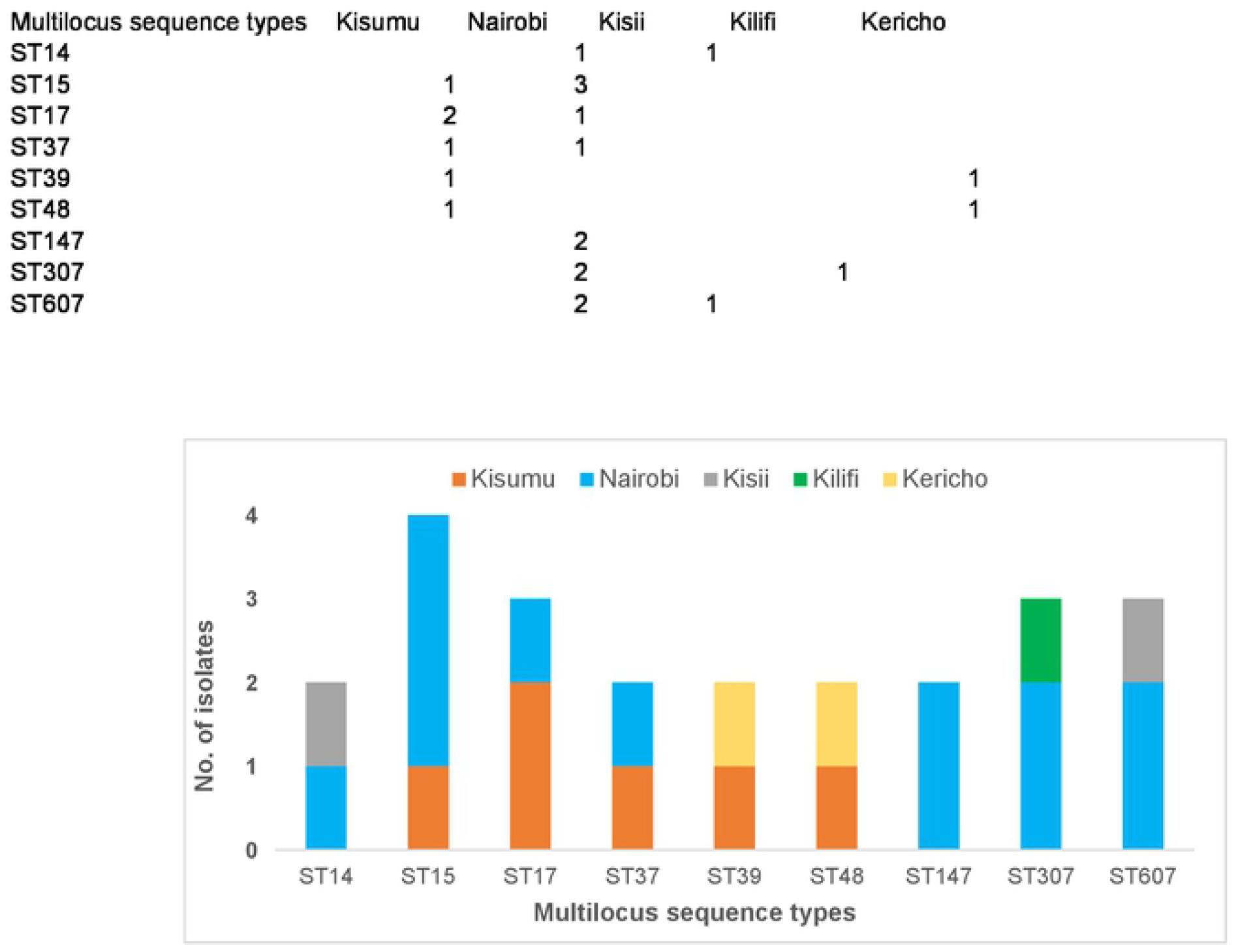

## Notes

### Competing Interest Statement

The authors have declared no competing interest.

